# PGC-1α isoforms coordinate to balance hepatic metabolism and apoptosis in inflammatory environments

**DOI:** 10.1101/703678

**Authors:** Mélissa Léveillé, Aurèle Besse-Patin, Nathalie Jouvet, Aysim Gunes, Stewart Jeromson, Naveen P. Khan, Sarah Sczelecki, Cindy Baldwin, Annie Dumouchel, Jorge Correia, Paulo Jannig, Jonathan Boulais, Jorge L. Ruas, Jennifer L. Estall

## Abstract

Liver is regularly exposed to changing metabolic and inflammatory environments. It must sense and adapt to metabolic need while balancing resources required to protect itself from insult. PGC-1α is a transcriptional coactivator that both coordinates metabolic adaptation to diverse stimuli and protects against inflammation. However, it is not known how PGC-1α integrates extracellular signals to balance metabolic and anti-inflammatory outcomes. PGC-1α exists as multiple, alternatively spliced variants expressed from different promoters. Primary mouse hepatocytes were used to evaluate the role(s) of different PGC-1α proteins in regulating hepatic metabolism and inflammatory signaling downstream of TNFα. PGC-1α1 and PGC-1α4 were expressed in hepatocytes and expression analysis uncovered shared and isoform-specific roles for these variants linked to metabolism and inflammation. PGC-1α1 primarily impacted gene programs of nutrient and mitochondrial metabolism, while TNFα signaling revealed that PGC-1α4 influenced several pathways related to innate immunity and cell death. Gain- and loss-of-function models illustrated that PGC-1α4 uniquely enhanced expression of anti-apoptotic gene programs and attenuated hepatocyte apoptosis in response to TNFα or LPS. This was in contrast to PGC-1α1, which reduced the expression of a wide inflammatory gene network, but did not prevent liver cell death in response to the cytokine. We conclude that PGC-1α variants have distinct, yet complimentary roles in hepatic responses to metabolism and inflammation and identify PGC-1α4 as an important mitigator of apoptosis.

## INTRODUCTION

The unique anatomical architecture of the liver allows it to perform a broad range of metabolic functions, but at the same time it exerts powerful immunocompetence, surveilling portal blood and acting as a protective barrier [1]. The liver must adapt quickly to various metabolic and inflammatory signals from the digestive tract or systemic circulation, while at the same time responding to changing glucose, metabolite and lipid homeostasis. Importantly, hepatic metabolism can be reprogramed by an inflammatory response [2], allowing a trade-off between energy destined for nutrient metabolism versus tolerance to infection. However, mechanisms helping to balance responses to metabolic demand with inflammation are not clear.

The peroxisome proliferator activated receptor gamma coactivator-1 alpha (PGC-1α) regulates many transcriptional programs related to nutrient metabolism, energy homeostasis and mitochondrial respiration [3] by binding to nuclear receptors and other transcription factors to enhance their activity [4]. PGC-1α also activates expression of gene programs within a broader set of biological functions in muscle [5–8] and liver [9–14].

Evidence suggests that PGC-1α is also an essential component of the inflammatory response. Over-expression in muscle protects mice from disease, exercise, and age-related inflammatory damage [15–18] and preservation of PGC-1α activity blunts lipopolysaccharide (LPS)-induced inflammatory damage to heart and kidney [19, 20]. Consistently, a low level of PGC-1α increases pro-inflammatory cytokine expression and inflammatory damage to muscle and liver tissue in response to cellular stress [17, 21, 22]. Over-expression of PGC-1α decreases expression of pro-inflammatory cytokines, while simultaneously inducing expression of secreted anti-inflammatory factors [17, 23].

Mechanistic understanding of links between inflammatory signaling and PGC-1α activity remain limited. Although PGC-1α is generally considered a coactivator, data suggest that PGC-1α indirectly represses NF-κB target gene transcription though coactivation of anti-inflammatory transcriptional networks linked to PPARs [18]. It may also bind directly to the p65 subunit of nuclear factor kappa-light-chain-enhancer of activated B cells (NF-κB) [24]. The primary function of PGC-1α is to increase the number and efficiency of mitochondria, an organelle important for energy production, but also for responding to extra- and intra-cellular signals to coordinate metabolic, inflammatory and cytotoxic responses[2, 25]. In this study, we show that differentially spliced variants of the PGC-1α protein have unique functions in regulating hepatocyte responses to concurrently integrate metabolic and inflammatory signals.

## RESULTS

### PGC-1α1 and PGC-1α4 are expressed in primary mouse hepatocytes

PGC-1α mRNA expression is controlled by different promoters (proximal and alternative) induced in a stimulus- and context-dependent manner [26–32]. This leads to expression of multiple splice variants of PGC-1α whose expression pattern is tissue-specific [33–37]. Due to high homology, splice variants of this protein cannot be fully distinguished by qPCR [38]. Using variant-specific non-quantitative PCR, we detected transcripts for *NT-PGC-1α-a* and *PGC-1α1* of the proximal promoter, and *PGC-1α-b* and *PGC-1α4* of the alternative promoter, when primary mouse hepatocytes were treated with the adenylyl cyclase activator, forskolin (Fk) activating both promoters [26] (Supplementary Fig. 1). PGC-1α-b has an alternative first exon, but is believed to regulate mitochondrial respiration similar to canonical PGC-1α1 [39]. PGC-1α4 undergoes splicing similar to NT-PGC-1α-a to create a truncated protein, but these two forms have different first exons. NT-PGC-1α-a also has over-lapping functions with PGC-1α1 in muscle and brown fat cells in respect to mitochondrial metabolism [31, 40, 41], while PGC-1α4 uniquely regulates myocyte hypertrophy [35]. There are no known functions for any of the alternative isoforms in liver. As full length PGC-1α1 can modulate tissue responses to inflammation, we sought to determine whether PGC-1α4 also influences inflammatory signaling and determine whether there are functional consequences of this within liver cells.

### PGC-1α1 and PGC-1α4 have distinct roles in the hepatic response to TNFα

We first compared the transcriptome of primary mouse hepatocytes by microarray following over-expression of PGC-1α1, PGC-1α4, or vector alone in the absence or presence of the inflammatory cytokine TNFα, a cytokine commonly associated with hepatic inflammation (Supplementary Fig. 2). More than 1000 genes changed by ≥2-fold following PGC-1α1 over-expression compared to vector alone (adj. p-value < 0.01), while only 24 were changed by PGC-1α4 and only 4 genes overlapped between the two lists (Fig 1A, Supplementary File 1). Following TNFα treatment, >4500 genes were changed ≥2-fold in hepatocytes over-expressing PGC-1α1 and >3000 for PGC-1α4, with 36% of the genes shared between isoforms (Fig 1A, Supplementary File 2). Clustering of PGC-1α4-modulated genes and comparison to levels in vector- or PGC-1α1-expressing hepatocytes suggested that the activity of PGC-1α4 relied heavily on the presence of TNFα (Fig 1B). Within this inflammatory context, PGC-1α4 had both over-lapping and distinct activity from PGC-1α1. Of the 2051 genes shared by the variants in TNFα-treated cells, the majority (91.5%) were regulated in the same manner (positively or negatively, Supplementary File 2). However, there were notable differences in some effector pathways for the two variants.

**Figure 1:**
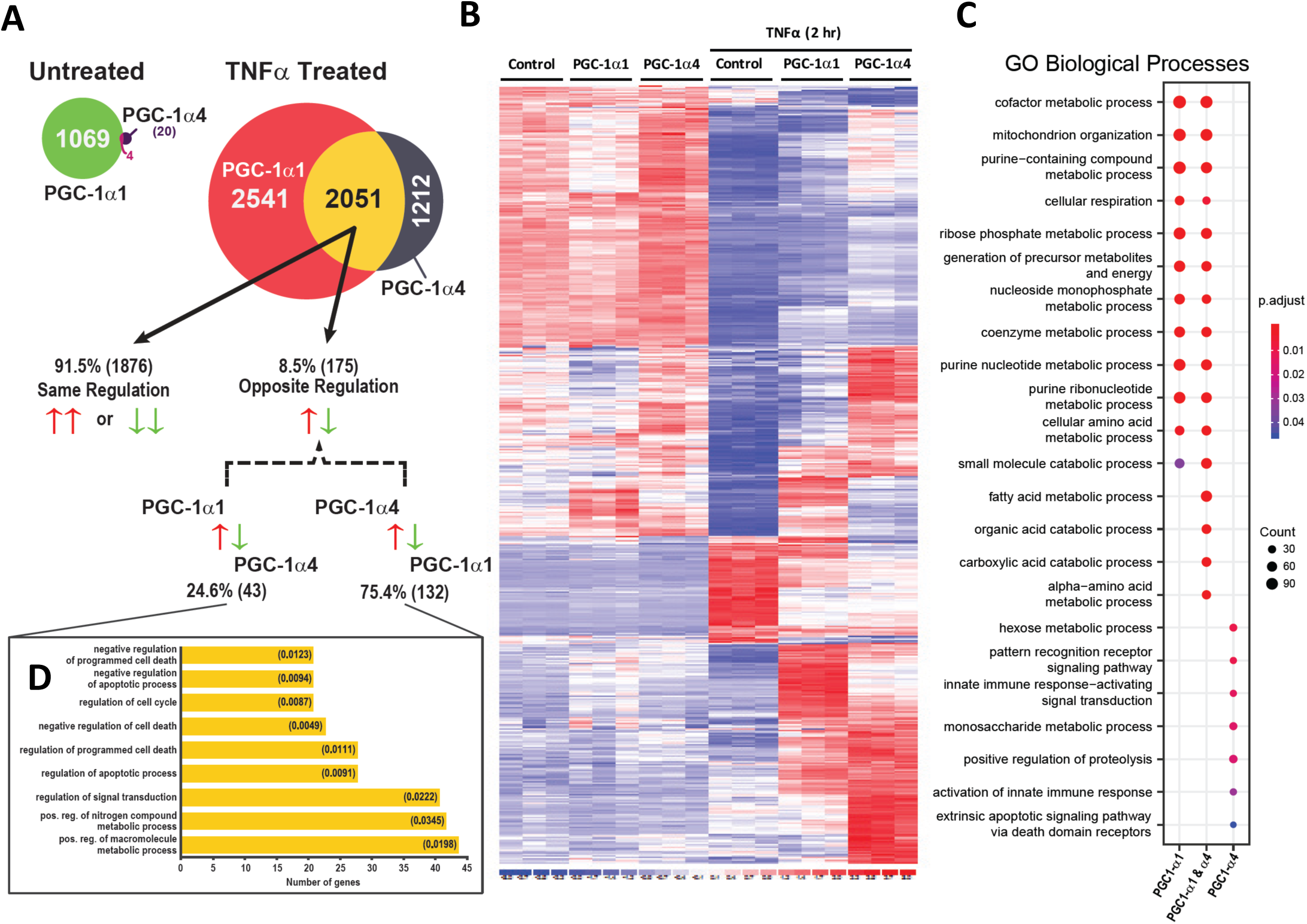
PGC-1α isoforms differentially regulate inflammatory and metabolic signaling pathways downstream of TNFα. Gene expression microarrays of mRNA isolated from primary mouse hepatocytes over-expressing either PGC-1α1, PGC-1α4, or vector control by adenoviral infection. A) Number of genes changed greater than 2-fold 48 hr following transduction in the absence or presence of 2 ng/mL TNFα (2 hr) (n = 3 biological replicates, adj. p-value <0.01). B) Clustering of genes significantly changed by over-expression of PGC-1α4 in primary hepatocytes in the presence of TNFα. C) Top 10 GO biological processes (adj. p-value <0.05) were identified from each list generated from TNFα-treated samples in A and listed on x-axis. Size of dot represents number of genes identified in each pathway, in comparison to other genotypes. D) GO biological processes (adj. p-value <0.05) associated with 175 genes regulated in the opposite direction. Data sets were generated using biological replicates (n=3) of each condition from one experiment.

To gain a global impression of biological process regulated by the PGC-1α variants in hepatocytes, we performed gene set enrichment analysis (GSEA). Gene sets relating to mitochondrial respiration and substrate metabolism were statistically enriched by both PGC-1α1 and PGC-1α4. PGC-1α1 predominantly regulated mitochondrial respiration, and glucose, amino acid and fatty acid metabolism, regardless of TNFα treatment (Supplementary File 3). This is consistent with known roles of PGC-1α1 in mitochondrial metabolism and supported by qPCR (Supplementary Fig 3A-D). Although we saw no effect of PGC-1α4 on these PGC-1α1 target genes, PGC-1α1 and PGC-1α4 shared many overlapping gene sets (Supplementary File 3). GSEA for PGC-1α4 in untreated hepatocytes centered on lipid metabolism (fatty acids and triglycerides), sterol metabolism and mitochondrial respiration, but individual gene effects were mild and most did not reach the 2-fold cut-off. However, with TNFα present, PGC-1α4-enriched pathways included regulation of transcription factor transport to the nucleus, innate immunity, responses to interferon/PAMP, TLR signaling, acute inflammation, and apoptosis. Overall, TNFα signaling revealed isoform-specific responses and highlighted processes related to the innate immune response and cell death unique to PGC-1α4.

To explore differential effects of the isoforms on inflammation, we performed gene ontology (GO) analysis on gene changes occurring only in the presence of TNFα. Top 10 GO pathways unique to each variant, or shared, are shown in Fig 1C. All of the top PGC-1α1-regulated processes focused on energy metabolism and were shared with PGC-1α4. However, 6 of the top pathways for PGC-1α4 were unique to this variant, including 6-carbon metabolism, proteolysis, immune signaling in response to pathogens, and regulation of cell death (Fig 1C). Interestingly, GO terms associated with the 175 shared genes regulated in an *opposite* manner by the variants (Supplementary File 4) centered mainly on cell death and apoptosis (Fig 1D). These data suggest that apoptosis is an important effector pathway *differentially* regulated by these two PGC-1α protein variants.

### PGC-1α1 and PGC-1α4 differentially regulate TNFα-induced hepatocyte apoptosis

Searching the microarray for candidate anti-apoptotic genes downstream of PGC-1α4, we identified *Birc2* (*Ciap1*) and *Tnfaip3* (also known as A20) (Fig. 1A), two anti-apoptotic proteins that prevent cell death downstream of inflammatory signaling. In a separate experiment, we confirmed their transcript levels were significantly higher in mouse primary hepatocytes over-expressing PGC-1α4 only in the presence of TNFα (Fig 2A). Related *Birc3* (*Ciap2*) was also increased by TNFα/PGC-1α4, while *Birc5* (Survivin) expression did not change. In addition, transcripts for apoptosis inhibitors *Naip* and *Xiap* were significantly increased by PGC-1α4, regardless of TNFα treatment. In contrast, over-expression of PGC-1α1 decreased expression of *Birc3, Birc5*, and *Tnfαip3* and had no effect on *Naip* and *Xiap* (Fig 2A). This data confirmed array and GO analysis (Fig 2D) illustrating that PGC-1α1 and PGC-1α4 have differential effects on apoptotic gene programs in response to TNFα. The predicted consequences of this were then demonstrated in primary mouse hepatocytes, where over-expression of PGC-1α1 increased cleaved caspase 3 (Fig 2B) and nucleosome fragmentation (Fig 2C) in response to TNFα treatment, while over-expression of PGC-1α4 almost completely blocked apoptosis compared to vector and PGC-1α1 (Fig 2B,C).

**Figure 2:**
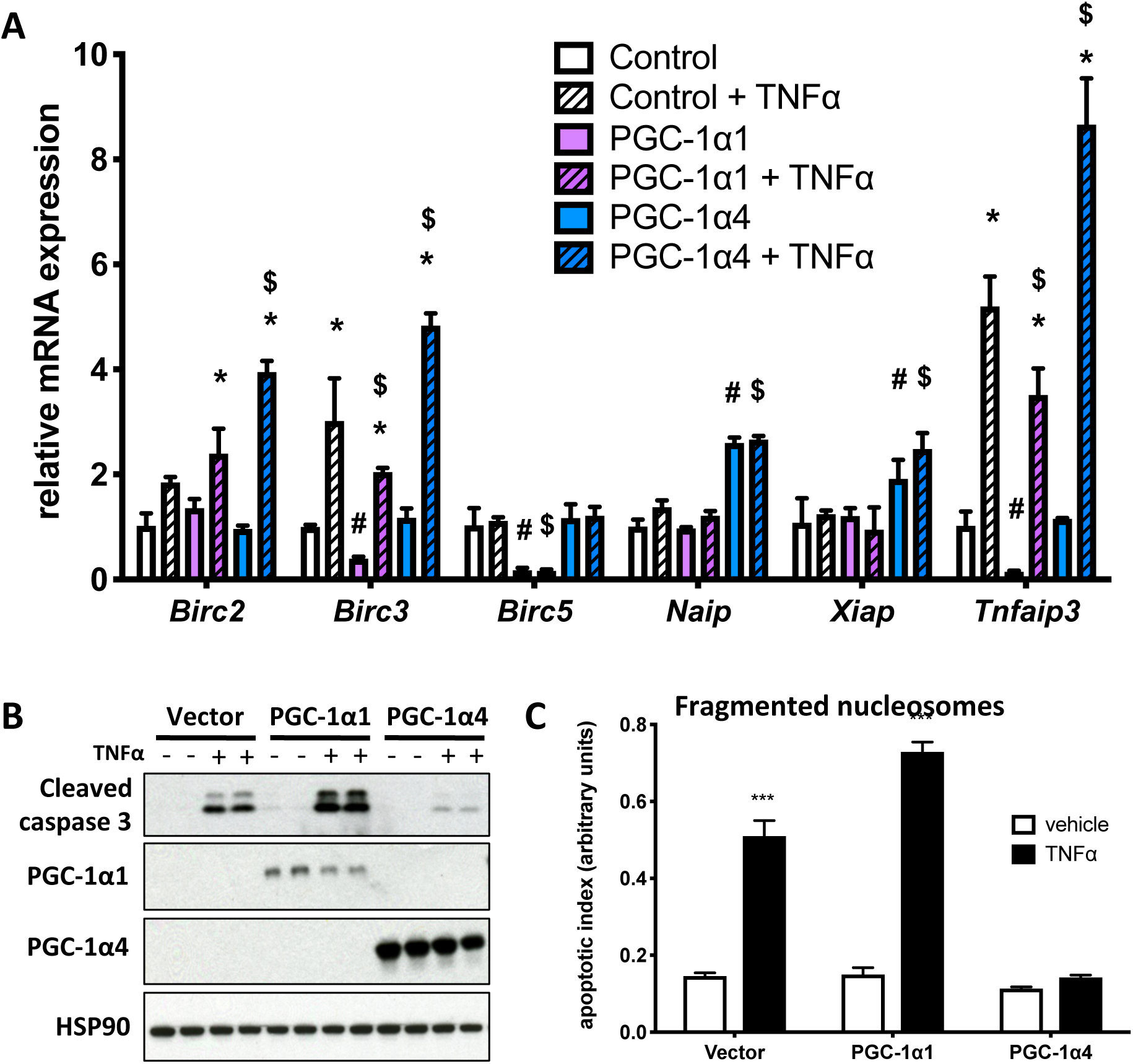
Over-expression of PGC-1α4 attenuates liver cell apoptosis induced by inflammatory signals. A) mRNA expression of primary mouse hepatocytes over-expressing PGC-1α1, PGC-1α4 or vector alone following 2-hr treatment with 2 ng/mL TNFα or vehicle (n=3). *p<0.05 effect of TNFα within each genotype. **^#^**p<0.05 TNFα response compared to Control + TNFα. **^$^**p<0.05 TNFα response compared to PGC-1α1 + TNFα. Data are biological replicates representative of 2-3 independent experiments. B) Western blot (n=2) and C) fragmented nucleosomes (n=4) in primary mouse hepatocytes over-expressing either PGC-1α1, PGC-1α4, or vector control by adenoviral infection, treated with or without 20 ng/mL TNFα for 8 hours. ***p<0.001 versus vehicle. Data are representative of 3 independent experiments.

**Figure 3:**
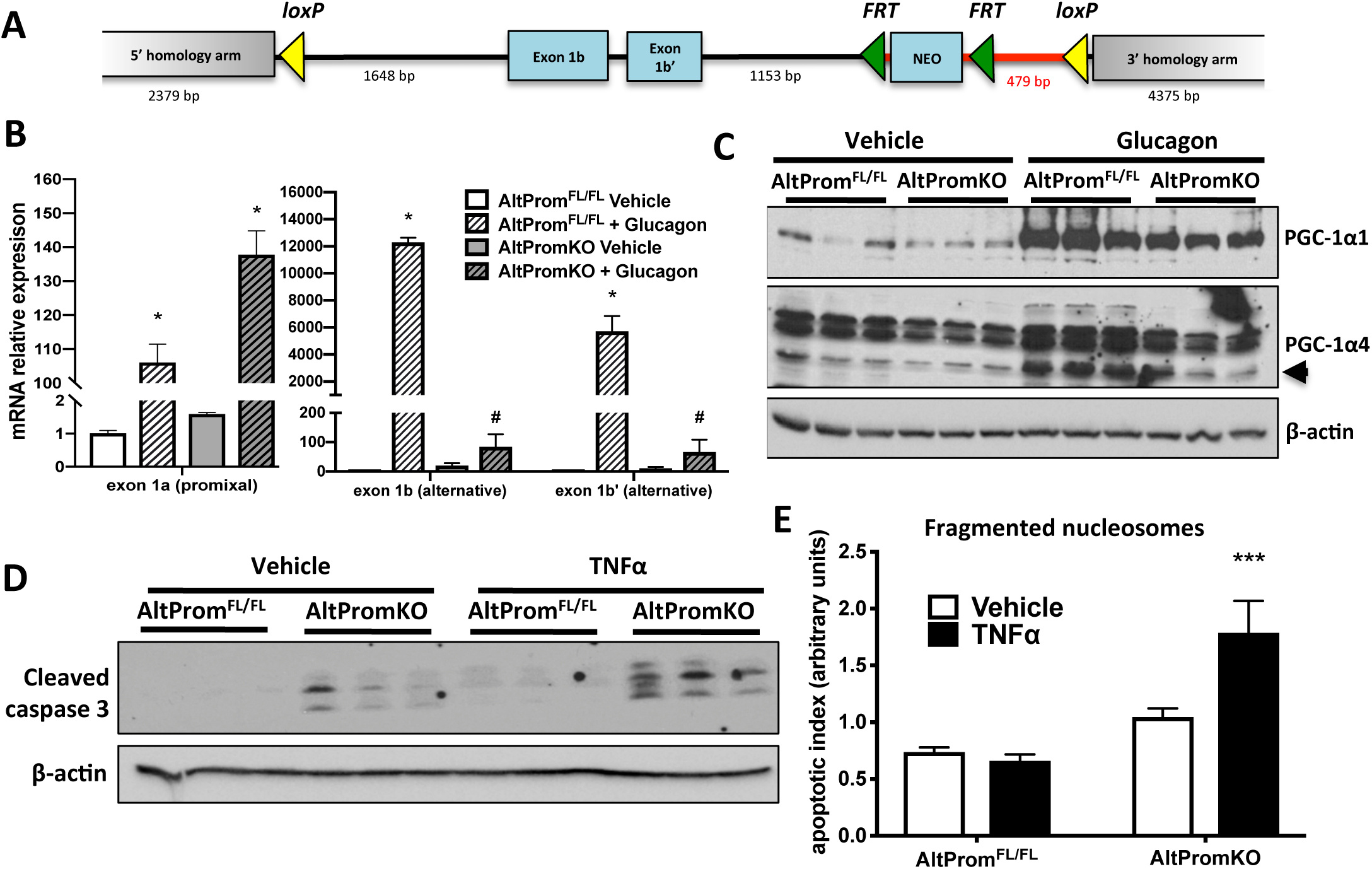
Loss of PGC-1α4 expression enhances apoptosis in response to TNFα. A) Targeting construct for creation of mouse allowing tissue-specific ablation of the alternative *Ppargc1a* promoter (AltProm^FL/FL^). B) mRNA of proximal and alternative *Pgc-1*α transcripts and C) western blot of proteins from primary mouse hepatocytes treated with 50 nM glucagon or vehicle. *p<0.05 versus AltProm^FL/FL^ Vehicle. #p<0.05 versus AltPromKO Vehicle, n = 3. D) Western blot and E) fragmented nucleosomes from primary mouse hepatocytes treated with 20 ng/mL TNFα or vehicle for 8 hours. ***p<0.001 versus AltProm^FL/FL^ Vehicle, n = 3. Bars are mean ± SEM of biological replicates in one experiment. Data are representative of 2 independent experiments.

We next investigated whether PGC-1α4 played an essential role in the hepatic apoptotic response. The commonly used model PGC-1α floxed mouse [22] ablates all *Ppargc1a* transcripts from both promoters, making it impossible to discern roles for any specific isoform. Thus, we created a model where only the alternative promoter was disrupted (AltProm^FL/FL^), blunting expression of variants containing exon 1b and 1b’ (including PGC-1α4), but not PGC-1α1 (Fig 3A). Efficiency of the promoter knockout was confirmed using glucagon, which significantly induced expression of PGC-1α transcripts (Fig 3B) and proteins (Fig 3C) from both the proximal and alternative promoter in control (AltProm^FL/FL^) cells. In contrast, ablation of the alternative promoter by crossing floxed mice with Alb-Cre^Tg^ mice (AltPromKO) blunted induction of transcripts only from the upstream promoter, with increases in proximal transcripts being similar to (or even higher than) control cells (Fig 3B). The 37kD PGC-1α protein induced by glucagon was almost completely ablated by knockout of the alternative promoter, identifying PGC-1α4 as the predominant truncated PGC-1α variant responsive to glucagon (Fig 3C). Consistent with PGC-1α4 being an important mediator of apoptosis, hepatocytes from AlbPromKO mice had higher basal and TNFα-induced cleaved caspase 3 levels (Fig 3D) and increased fragmented nucleosomes in response to the cytokine (Fig 3E) compared to cells from littermate controls. Taken together with gain-of-function data (Fig 2), PGC-1α4 appears to have the unique ability to prevent inflammatory-mediated apoptosis in liver cells.

### TNFα signaling influences localization of PGC-1α4 within liver cells

It was curious to us why the activity of PGC-1α4 was dependent on activation of TNFα signaling. We first investigated whether TNFα itself regulates transcription from either *Ppargc1a* promoter, on its own or in conjunction with other signals. Unlike glucagon treatment, which activates cAMP:CREB and acutely induced both promoters, TNFα treatment did not significantly influence levels of PGC-1α transcripts derived from proximal (PGC-1α1 or NT-PGC-1α-a) or alternative (PGC-1α-b or PGC-1α4) promoters over a 24-hour period (Fig 4A,B). TNFα also had no impact on the capacity of glucagon to induce either promoter. Since TNFα treatment substantially increased the number of PGC-1α4 gene targets in our microarray (Fig 1A), we proposed that inflammatory signals might regulate the PGC-1α proteins, or their activity, in other ways. Using immunofluorescence, we noted that over-expressed PGC-1α4 localized almost exclusively to the cytoplasm of liver cells, suggesting that nuclear exclusion prevents PGC-1α4 from regulating nuclear gene expression. Following addition of TNFα, a significant proportion of PGC-1α4 was detected in the perinuclear and nuclear compartments (Fig 4C). Cell fractionation confirmed that PGC-1α4 protein was only detected in the nuclear pellet following TNFα treatment. In contrast, PGC-1α1 localized exclusively to the nucleus of liver cells regardless of treatment conditions (Fig 4D). PGC-1α4 migrated at a variety of molecular weights, but the highest molecular weight form was predominant in the nuclear fraction. Thus, nuclear exclusion may be a mechanism to maintain PGC-1α4 in an inactive state in the absence of inflammatory stimuli.

**Figure 4:**
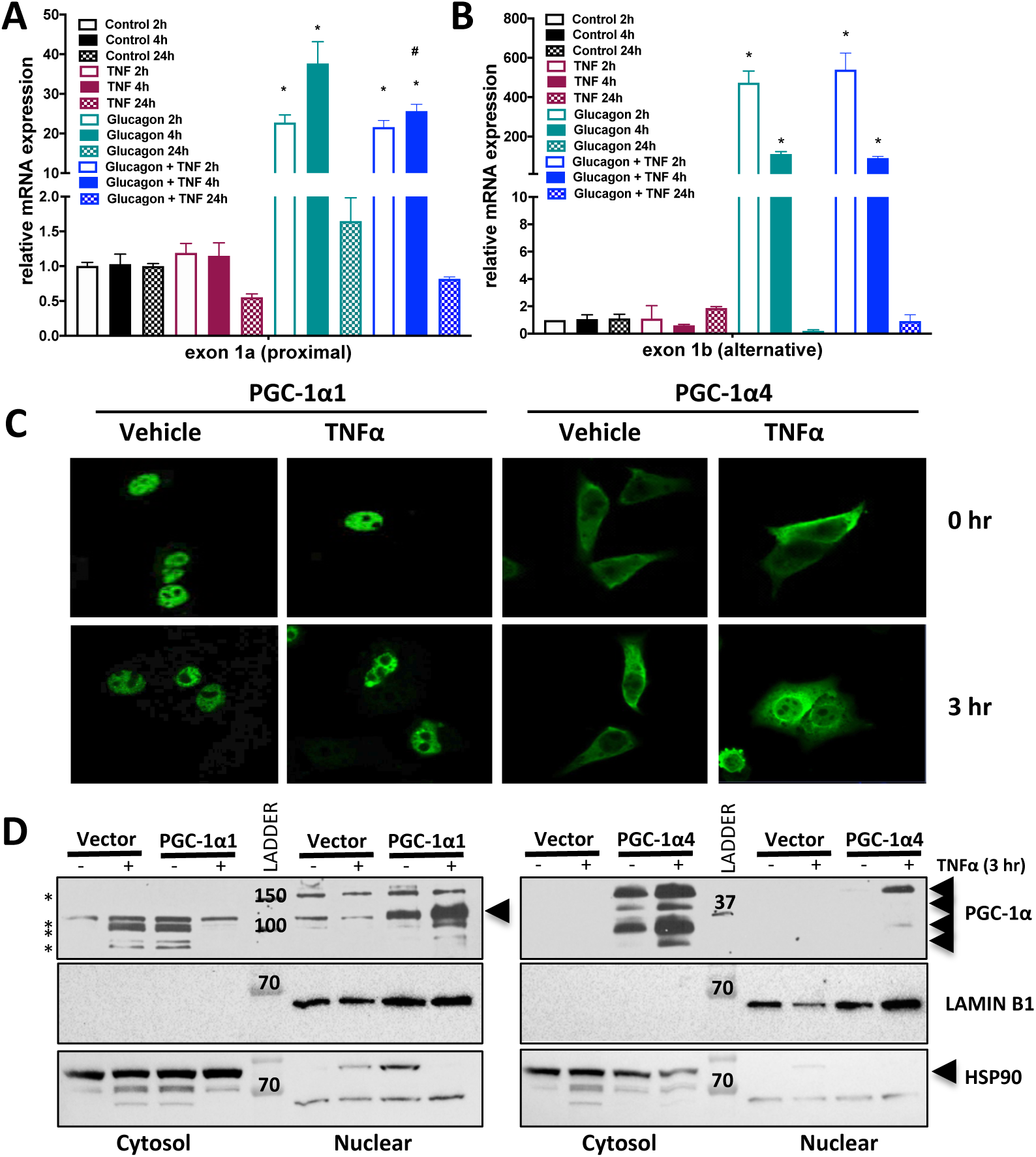
TNFα signaling promotes mobilization of cytoplasmic PGC-1α4 to nuclear and perinuclear regions. A,B) mRNA levels of transcripts expressing either exon 1a of the proximal promoter, or exon 1b of the alternative promoter from primary mouse hepatocytes treated with control (PBS), glucagon (50 nM) or TNFα (20 ng/mL) for indicated times. *p<0.05 TNF effect compared to control, #p<0.05 TNF effect compared to glucagon. C) Confocal imaging of H2.35 mouse liver cells transfected with plasmids expressing V5-tagged PGC-1α1 or PGC-1α4 treated with 50 ng/mL TNFα or vehicle (PBS) for 3 hours. D) Cell fractionation of H2.35 mouse liver cells transduced with adenovirus expressing control vector, PGC-1α1 or PGC-1α4 and treated with 50 ng/mL TNFα or vehicle (PBS) for 3 hours. *non-specific bands. Data are representative of 3 independent experiments.

### PGC-1α4 does not repress the pro-inflammatory gene program in liver cells

Since anti-apoptotic gene programs are often regulated by NF-κB, we hypothesized that PGC-1α1 and PGC-1α4 might have differential effects on NF-κB activity. Basal expression of a 3x NF-κB response element reporter was increased in primary hepatocytes when co-expressed with PGC-1α1 (Supplementary Fig 4A). However, consistent with previous findings [18, 24], TNFα-induced activity of the reporter was significantly blunted by high PGC-1α1 (Supplementary Fig 4A) as well as basal and/or TNFα-induced levels of pro-inflammatory genes *Mcp-1*, *Tnfα*, *Iκbα* and *Ccl5* in primary hepatocytes (Supplementary Fig 4B). This confirmed a strong inhibitory role for PGC-1α1 on NF-κB signaling in liver cells. In contrast, PGC-1α4 had no effect on NF-κB reporter activity and little impact on target pro-inflammatory genes, except to potentiate the *Tnfα* response similar to anti-apoptotic targets (Supplementary Fig 4B).

In an attempt to identify transcription factors with links to apoptosis and cell death downstream of PGC-1α variants, we probed array data for transcription factor motifs enriched in genes: changed by PGC-1α1 or PGC-1α4 alone; oppositely regulated by the two variants; or shared when TNFα was present using iRegulon (Supplementary Table S4) and DiRE (Supplementary Fig 5). Many motifs not previously associated with PGC-1α were identified including: ETV6 (only PGC-1α1); SP4, the NFY complex, IRF6, GM7148, TGIF2, HSF4, and E2F1DP1 (only PGC-1α4); and IRF4, LK4, NR1H2 (LXRβ), ZBTB33 (KAISO), ZFP143, and PITX2 (shared). Among the 175 genes oppositely regulated by the variants, binding sites for STAT, SPIB, NFATC2, and KLF4 were found. ST18 (also known as MYT3 or NZF3), was enriched when we focused on the gene list implicated in cell survival (Fig 1D). Narrowing our focus to transcription factors implicated in cell death and inflammation, we evaluated whether over-expression of the PGC-1α variants modulated expression of their target genes. PGC-1α4 had no significant effects on mRNA expression of any target genes, while PGC-1α1 repressed SP4, NF-Y, and STAT targets (Supplementary Fig 6A,B,C), and increased IRF4 targets *Tnfrsf17* and *Nip3* (Supplementary Fig 6D). In summary, PGC-1α1 generally repressed NF-κB activity and the transcription of genes involved in inflammation and cell death. In contrast, PGC-1α4 differentially enhanced a distinct program of anti-apoptotic factors in hepatocytes only in the presence of TNFα-signaling, but this did not appear to be via enhancement of NF-κB.

**Figure 5:**
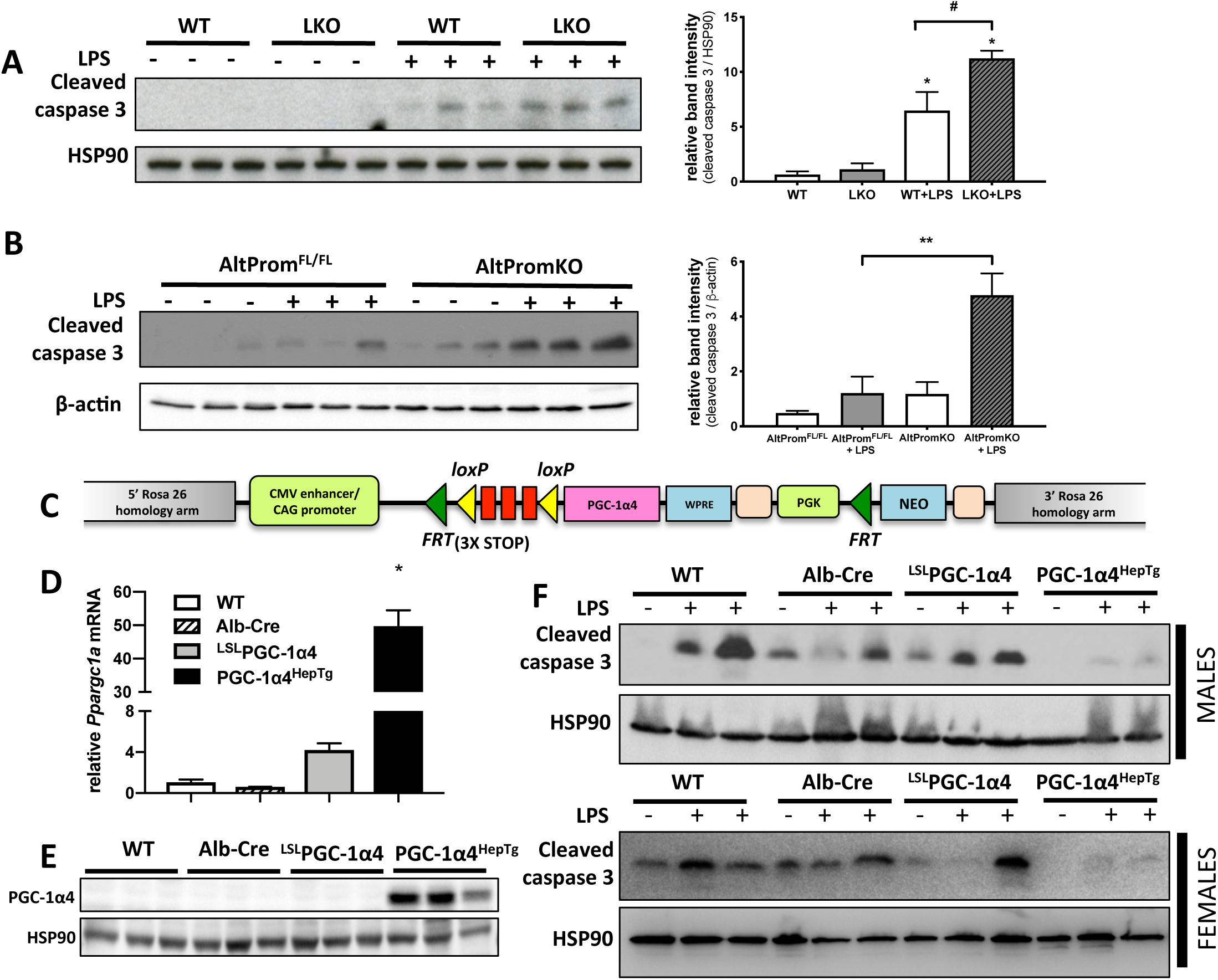
PGC-1α4 is necessary and sufficient to prevent LPS-induced liver cell apoptosis. A) Western blot of liver protein from male WT or LKO mice (n = 3 mice) 6 hours following injection of LPS (2 mg/kg) or vehicle (PBS). *p<0.05 versus WT control levels. # p<0.05 versus WT + LPS levels. B) Western blot of liver protein from mice 6 hours following tail-vein injection of 2 mg/kg LPS or vehicle (PBS) (n = 3). C) Targeting construct for transgenic mouse allowing tissue-specific over-expression of PGC-1α4. D) mRNA and E) protein from livers of mice (n = 3) following breeding of ^LSL^PGC-1α4 mice with Albumin-Cre^Tg^ mice to drive PGC-1α4 expression only in hepatocytes (PGC-1α4^HepTg)^. *p<0.05 versus WT control, n=3 mice. F) Western blot of liver protein from male and female mice 6 hours following tail-vein injection of 2 mg/kg LPS (n = 6 of each sex) or vehicle (PBS) (n = 2 of each sex).

### PGC-1α4 prevented lipopolysaccharide (LPS)-induced hepatocyte cell death in vivo

Lastly, we investigated whether PGC-1α4 was important for regulating liver cell apoptosis downstream of inflammatory signals *in vivo*. Mice lacking all isoforms of PGC-1α in liver had increased cleaved caspase 3 levels when exposed to LPS (Fig 5A). When we ablated transcription from the alternative *Ppargc1a* promoter only, protein extracts from livers of AltPromKO mice treated with LPS also had increased cleaved caspase 3 compared to littermate AltProm^FL/FL^ control livers (Fig 5B). To determine whether increases in PGC-1α4 would conversely prevent LPS-induced apoptosis, we created a novel ROSA26 knockin mouse with PGC-1α4 cDNA positioned downstream of a Lox-Stop-Lox cassette (Fig 5C). There was a small increase in PGC-1α4 transcripts in the absence of Cre-recombinase (^LSL^PGC-1α4) suggesting a low level of leaky transgene expression. However, upon crossing with Albumin-Cre^Tg^ mice, PGC-1α4^HepTg^ mice over-expressed PGC-1α4 only in liver cells ∼50-fold higher than controls (Fig 5D,E). Supporting the anti-apoptotic role for hepatic PGC-1α4, there were reduced levels of cleaved caspase 3 in livers of both male and female PGC-1α4^HepTg^ mice following injection of LPS (Fig 5F).

## DISCUSSION

In the current study, we found that various non-canonical PGC-1α protein variants are expressed in murine liver and that these differentially regulate metabolism and hepatic inflammatory signaling. Gene set enrichment analysis revealed that in the presence of the inflammatory cytokine TNFα, PGC-1α4 influences innate immunity and cell death, while PGC-1α1 remains primarily associated with mitochondrial function and metabolic processes. Gene ontology (GO) analysis and qPCR illustrated that genes implicated in cell death and apoptosis are often oppositely regulated by these two variants. In primary liver cells, PGC-1α4 has the unique ability to blunt apoptosis in response to TNFα, a function that may be controlled by shuttling of PGC-1α4 between cytoplasm and nucleus. We conclude that alternative forms of PGC-1α are inducible and co-expressed in liver, giving rise to isoforms that coordinate the cellular response to inflammatory signaling. These findings give mechanistic insight into how PGC-1α, as a family of proteins, facilitate parallel adaptation to metabolic demand and mitigation of inflammatory damage in cells.

Immune responses to danger signals are metabolically challenging and can lead to a trade-off between maintaining highly energy demanding processes of nutrient metabolism versus adaptation to inflammatory stimuli [2]. Inflammation itself often inhibits metabolism and impedes mitochondrial function. Here, we show that signaling via TNFα or LPS leads to a shift in the PGC-1α gene program downstream of PGC-1α1 and PGC-1α4, ensuring that concurrent inflammatory signaling does not impede the ability of liver cells to respond to metabolic need. This mechanism represents an additional layer by which the family of PGC-1α proteins help balance an integrated metabolic response that is modulated by the inflammatory status of the liver.

It is now well established that PGC-1α is a family of proteins created by alternative splicing of the *PPARGC1A* gene in many tissues including skeletal muscle [35, 36, 38], brown adipose tissue [37, 42], and liver [33]. However, a functional role for many of the alternative isoforms remains unknown. While some PGC-1α variants share overlapping functions with canonical PGC-1α1 [30, 31, 34, 39, 43], PGC-1α4 has distinct effector pathways in muscle and brown adipose tissue [35, 44]. We show here that PGC-1α1 and PGC-1α4 have differential effects on cell death downstream of inflammatory signals. PGC-1α4 almost completely blocks apoptosis *in vitro* and *in vivo*. While PGC-1α1 decreases expression of a broad program of inflammatory genes, it did not inhibit cell death in response to TNFα in hepatocytes. In fact, PGC-1α1 has been shown to induce apoptosis through PPARγ, TFAM, generation of reactive oxygen species, or Ca^2+^ signaling [45–49], but may also attenuate cell death by activation of p38/GSK3B/Nrf-2 axis or p53 [50, 51]. Newly identified PGC-1α splice variants from differentially regulated promoters add a layer of complexity, but also may explain conflicting data in previous reports.

An obvious candidate effector in inflammation-mediated apoptosis is NF-κB. Consistent with previous studies [52, 53], we show that PGC-1α1 represses NF-κB activity. However, unlike PGC-1α1, our evidence suggests no impact of PGC-1α4 on this transcription factor. Although PGC-1α4 shares the complete activation domain of PGC-1α1, its alternative exon 1 and significant C-terminal truncation may lead PGC-1α4 to regulate a different set of DNA-binding proteins. Our microarray identifies multiple pathways differentially regulated by the two variants, including those targeted by NF-κB, SP4, NF-Y, STAT and IRF4. However, in our model system, PGC-1α4 did not behave as a traditional transcriptional coregulator for many of their gene targets. One possible explanation could be that PGC-1α4 instead promotes novel splicing events to create alternative gene products, similar to the function of related PGC-1α2 and PGC-1α3 variants [54]. Aberrant alternative splicing can substantially affect cellular function and is associated with disease. For example, alternative splicing of TNFα-regulated genes (such as *Tnfaip3/A20*) produces protein variants with distinct roles in cell death and cell survival [55].

While the exclusive nuclear localization of PGC-1α1 supports its function as a transcriptional coactivator, the ability of PGC-1α4 to shuttle between compartments suggests that it might interact with transcription factors in the cytoplasm and/or regulate their entry into the nucleus, a possibility also supported by our GSEA analysis. Interferon (INF) regulatory factors (IRFs) are well-known transcription factors that shuttle in response to inflammatory stimuli [56] and our data suggest that both PGC-1α1 and PGC-1α4 converge on interferon signaling in liver cells. Canonical PGC-1α1 has been associated with interferon response in the contexts of HCV infection and thermogenesis [57, 58]. Interestingly, three interferon regulatory factors (IRF1, IRF4, IRF6) were identified in our motif enrichment analysis and numerous studies implicate interferons as critical regulators of apoptosis [59, 60]. Although we focused on TNF signaling, our data suggest that PGC-1α1 and PGC-1α4 might also regulate the interferon response; however, further studies are necessary to confirm this hypothesis. Overall, mechanisms underlying the anti-apoptotic role of hepatic PGC-1α4 appear complex, possibly involving interaction with cytoplasmic proteins, dominant-negative effects on other PGC-1α variants, or regulation of alternative splicing of genes implicated in apoptosis.

PGC-1α4 shares many similarities to another isoform, NT-PGC-1α, which is transcribed from the proximal promoter. Both have two N-terminal nuclear exclusion signals and three putative phosphorylation (S190, S237, and T252) sites, which regulate nuclear shuttling of NT-PGC-1α [42]. Our data are consistent with reports describing cytoplasmic to nuclear movement of other truncated variants of PGC-1α [37, 42]. Given similarities between these proteins, it is possible that NT-PGC-1α localization is also regulated by inflammation similar to PGC-1α4, and while likely, it remains to be seen whether PGC-1α4 and NT-PGC-1α have overlapping functions in terms of inflammation and apoptosis.

In conclusion, coordinated activity of PGC-1α isoforms allows fine-tuning of metabolic and inflammatory networks, supporting efficient adaptation to energy demand within the highly dynamic and inflammatory environment of the liver. Our data imply that boosting expression of specific PGC-1α isoforms could allow liver cells to more efficiently respond to energy demand when faced with both high metabolic and inflammatory challenges associated with metabolic disease. Offsetting this balance could result in inefficient nutrient metabolism and/or inappropriate responses to inflammatory stimuli, which may play a role in inflammatory liver diseases such as non-alcoholic steatohepatitis (NASH) or cirrhosis.

## MATERIALS AND METHODS

### Mice

Hepatocyte-specific PGC-1α knockout mice (LKO: *Ppargc1a*^fl/fl, Alb-cre^) were generated as previously described [10, 11, 22]. Age-matched, male mice on a C57BL/6J background were used. Tissue-specific PGC-1α4 over-expressing mouse line (^LSL^PGC-1α4) was generated by inserting PGC-1α4 cDNA downstream of a Lox-stop-Lox cassette at the ROSA26 locus (Supplementary Fig. 7A). *Ppargc1a* Alternative Promoter Knock-out (AltPromKO) mice were generated by inserting LoxP sites flanking exon 1b and 1b’ of the alternative *Ppargc1a* promoter (Supplementary Fig. 7B). Experiments were performed in accordance with IRCM institutional animal care and use committee regulations.

### Mouse housing, diets, and lipopolysaccharide treatment

Mice were maintained on *ad libitum* chow (Tekland #2918) at 22°C (12h light/dark cycle). For LPS treatment, livers of 10-week-old male or female mice were harvested 6 hours after tail-vein injection of LPS (2 mg/kg, Invivogen) or vehicle (PBS).

### Primary hepatocyte isolation and treatment

Primary mouse hepatocytes from 12-week-old mice were isolated and cultured as previously described [22]. One day after isolation, hepatocytes were infected with adenovirus (5 MOI) overnight and starved of insulin and dexamethasone for 24 hours prior to treatment with tumor necrosis factor alpha (TNFα) (Fitzgerald) at 2 ng/mL for 2 hours for signaling/gene expression, or 20 ng/mL for 8 hours for apoptosis. Apoptosis was measured by Cell Death Detection ELISA (Roche). For reporter assays, cells were transfected (Lipofectamine) with a construct expressing firefly luciferase downstream of 3x NF-κB response elements. Activity was normalized to total protein following quantification using the Dual Luciferase Reporter Assay System (Promega).

### Protein isolation, immunoprecipitation and immunoblotting

Proteins were solubilized in radioimmunoprecipitation assay buffer containing protease and phosphatase inhibitors. Hepatic PGC-1α was immunoprecipitated from liver using anti-PGC-1α (Millipore, ST1202) in 1% Triton/TBS. Elutes and total proteins were resolved by SDS-PAGE, blotted, and probed with antibodies (Supplementary Table 1).

### Immunofluorescence

H2.35 cells cultured in DMEM, supplemented with 10% Fetal Bovine Serum (FBS, Wisent), 1% penicillin/streptomycin, 0.2 μM dexamethasone were incubated on poly-L-lysine coated coverslips and transfected with V5-tagged PGC-1α variants for 24 hours (Lipofectamine). Cells were starved overnight of dexamethasone prior to TNFα treatment (50 ng/ml) for 3 hours and fixation with 4% paraformaldehyde. Triton-permeabilized cells were incubated with anti-V5 antibody overnight, followed by Alexa 488-conjugated secondary antibody to visualize proteins.

### Cell fractionation

H2.35 cells transduced with adenovirus expressing control vector, PGC-1α1 or PGC-1α4 were starved overnight of dexamethasone prior to TNFα treatment (50 ng/ml) for 3 hours. Cell pellets were washed in PBS and resuspended in Lysis Buffer (10 mM Hepes (pH 7.5), 10 mM KCl, 3 mM MgCl2, 0.35 M sucrose, 0.1% NP40, 3 mM 2-mercaptoethanol, 0.4 mM PMSF, 1 uM pepstatin A, 1 uM leupeptin and 5 ug/ml aprotinin). After centrifugation, supernatants were kept as cytoplasmic fraction. The pellet (nuclear fraction) was washed twice with lysis buffer, resuspended in Buffer A (3 mM EDTA, 0.2 mM EGTA, 1 mM dithiothreitol, 100 mM NaCl and 0.8% NP40) and sonicated for 10 minutes (cycles of 30 seconds ON and 30 seconds OFF). Equal amount of proteins were resolved by SDS-PAGE.

### Microarray and Gene set enrichment analysis

mRNA was isolated from primary mouse hepatocytes infected with adenovirus expressing PGC-1α1, PGC-1α4 or vector control treated with 2 ng/mL TNFα or vehicle (PBS) for 2 hours (n = 3) and gene expression profiles generated using Affymetrix Mouse Genome 430 2.0 Arrays. Raw CEL files were normalized using RMA [PMID: 12925520] and annotated using biomaRt [PMID: 16082012]. Raw data and sample annotation are available on GEO (GSE132458).

Gene set enrichment analysis was performed using javaGSEA software (version 3.0 – build: 01600) on chip data using the Gene Ontology processes (number of permutations = 1000, Permutation type = gene_set, Chip platform = Affy_430_2.0_mgi (version 2011) from the Mouse Genome Database. The *Ppargc1a* probe (1434099_at) was removed prior to analysis to eliminate over-expression bias. Full GSEA results are provided in Supplemental File 1. A MySQL database generated lists of genes significantly regulated (adj. p-value < 0.01, Log10 FC ≥ 0.3 or ≤ −0.3). Full lists are provided in Supplemental Files 2 (untreated samples) and 3 (TNFα–treated samples).

Clustering based on PGC-1α4-regulated genes was performed using dChip software. Over-representation analysis (ORA) of Gene Ontology processes was performed using ClusterProfiler and the mouse genome-wide annotation in R (www.r-project.org). The top 10 statistically over-represented processes were determined for each condition, merged in to one list, and represented as a dot plot (adj. p-value < 0.05, correction method = Bonferroni). For 175 genes regulated oppositely by the variants, ORA was performed using g:Profiler (adj. p-value < 0.05, correction method = g:SCS threshold). Gene lists were evaluated for enrichment of transcription factor signatures and binding sites in the proximal promoters and distant regulatory elements using iRegulon and DiRE (http://dire.code.org) with default analysis settings.

### RNA isolation, PCR, and quantitative RT-PCR

RNA was isolated from frozen tissue or cells using TRIzol (Invitrogen). 1 μg of RNA treated with DNase I was reverse-transcribed using the High Capacity Reverse Transcription Kit (Applied Biosystems). cDNA was quantified using SYBR Green PCR master mix (Bioline) and normalized to Hypoxanthine-guanine phosphoribosyltransferase (*Hprt*) mRNA using the ΔΔCt threshold cycle method. Presence or absence of PGC-1α variants was confirmed using isoform-specific primers by conventional PCR and sequencing (Supplementary Tables 2 and 3).

### Statistical analysis

Sample sizes were based on previous experience with assays and knowledge of expected variance. Normal distribution and homoscedasticity of data were tested by Shapiro−Wilks and Bartlett tests, respectively. Parametric tests were used if distributions normal and variances equal. One-way or Two-way analysis of variance were followed by Tukey’s (one-way) or Dunnett’s multiple comparisons (two-way) post-hoc test using GraphPad Prism software. Data are expressed as mean ± SEM unless otherwise indicated.

## ACKNOWLEDGEMENTS

We thank Dr Bruce Spiegelman for generating the AlbProm^FL/FL^ mouse line and members of the IRCM animal, microscopy, and molecular biology core facilities for invaluable technical assistance.

## Author contributions

ML, ABP, NJ, SJ, JLR and JLE designed concept and experiments. ML, ABP, NJ, SJ, NPK, SS, CB, AD, JC, JB and PJ performed and analyzed experiments. ML, ABP, NJ, SJ, JLR and JLE wrote the manuscript. All authors reviewed the manuscript.

---

**Supplementary Figure 1:**
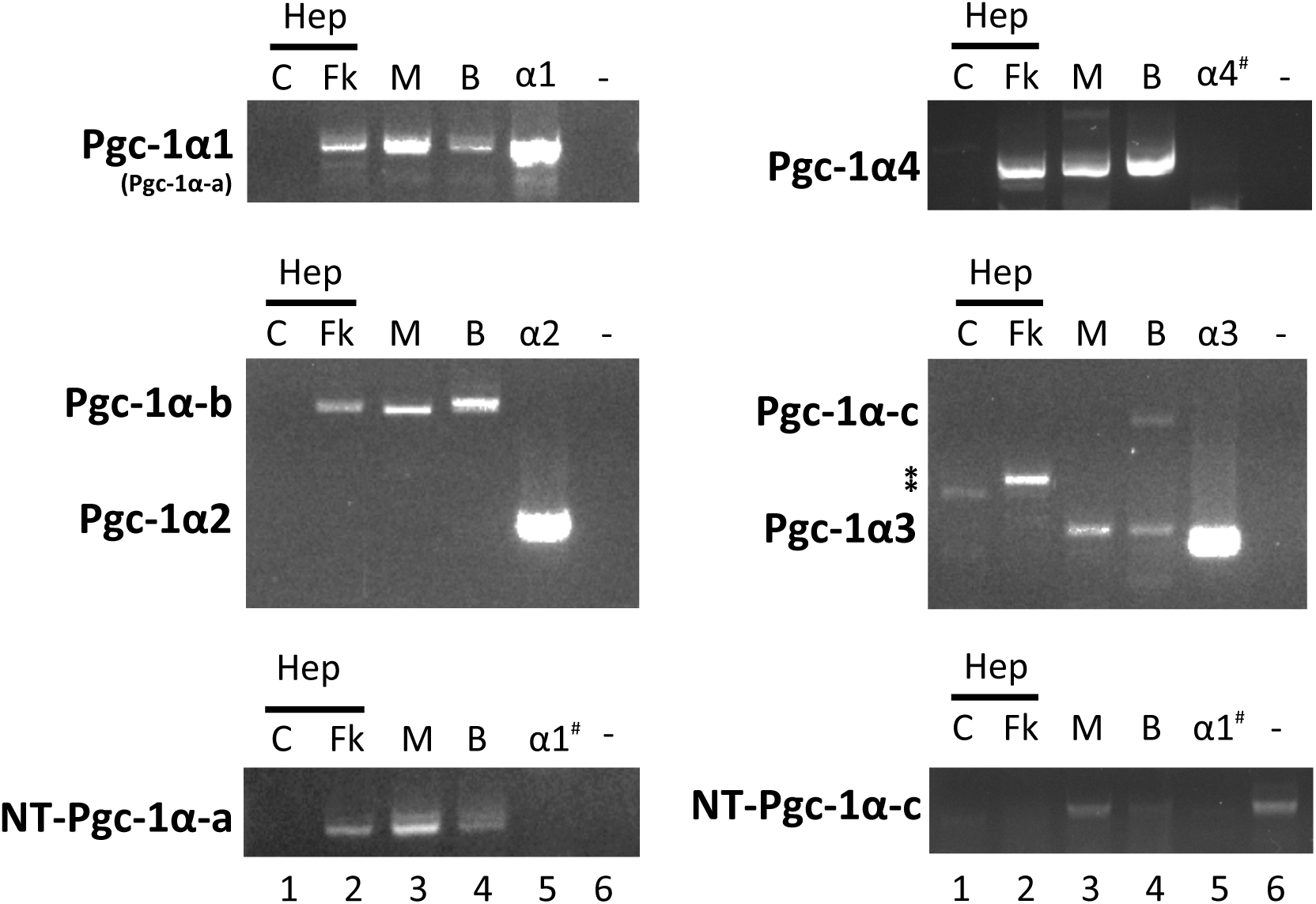
Figure 1: Expression levels of *Ppargc1a* transcripts in mouse tissues. Bands represent PCR products specific to each PGC-1α isoform, amplified from cDNA. Lanes 1 and 2: Primary mouse hepatocytes treated with 10 nM forskolin or control vehicle (DMSO) for 3 hours. C – vehicle control, Fk – forskolin treated. Lane 3: Mouse muscle (M), Lane 4: Mouse brown adipose tissue (B). Lane 5: Primary mouse hepatocytes over-expressing the indicated PGC-1α isoform (positive and negative controls). Control vectors for some variants were not available, in this case, # represents control performed on cells over-expressing a structurally similar variant to demonstrate specificity and non-cross-reactivity of primer sets. Lane 6: Water control.

**Supplementary Figure 2:**
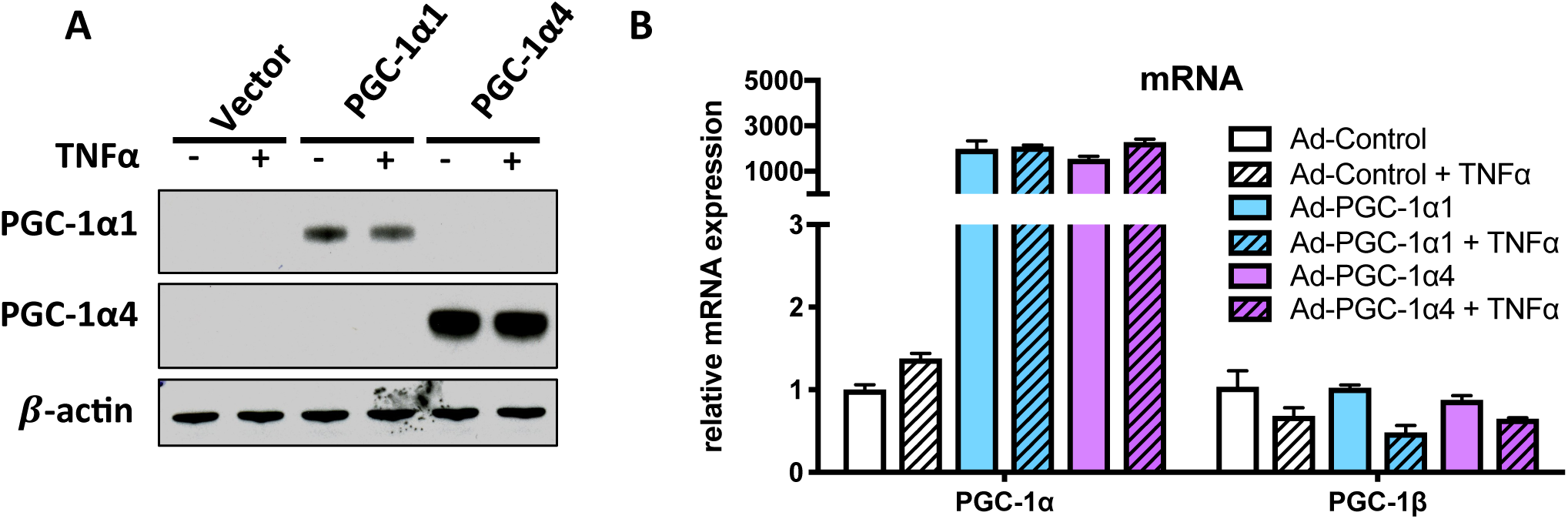
Relative levels of PGC-1 mRNA and protein following over-expression in primary mouse hepatocytes and TNFα treatment. **A**) Western blot of proteins and B) relative mRNA levels in primary hepatocytes 48 hours following transduction with an adenovirus expressing cDNA for PGC-1α1, PGC-1α4 or vector alone (Ad-CMV-GFP). Prior to harvest, cells were treated for 2 hours with 20 ng/mL TNFα or vehicle alone (PBS).

**Supplementary Figure 3:**
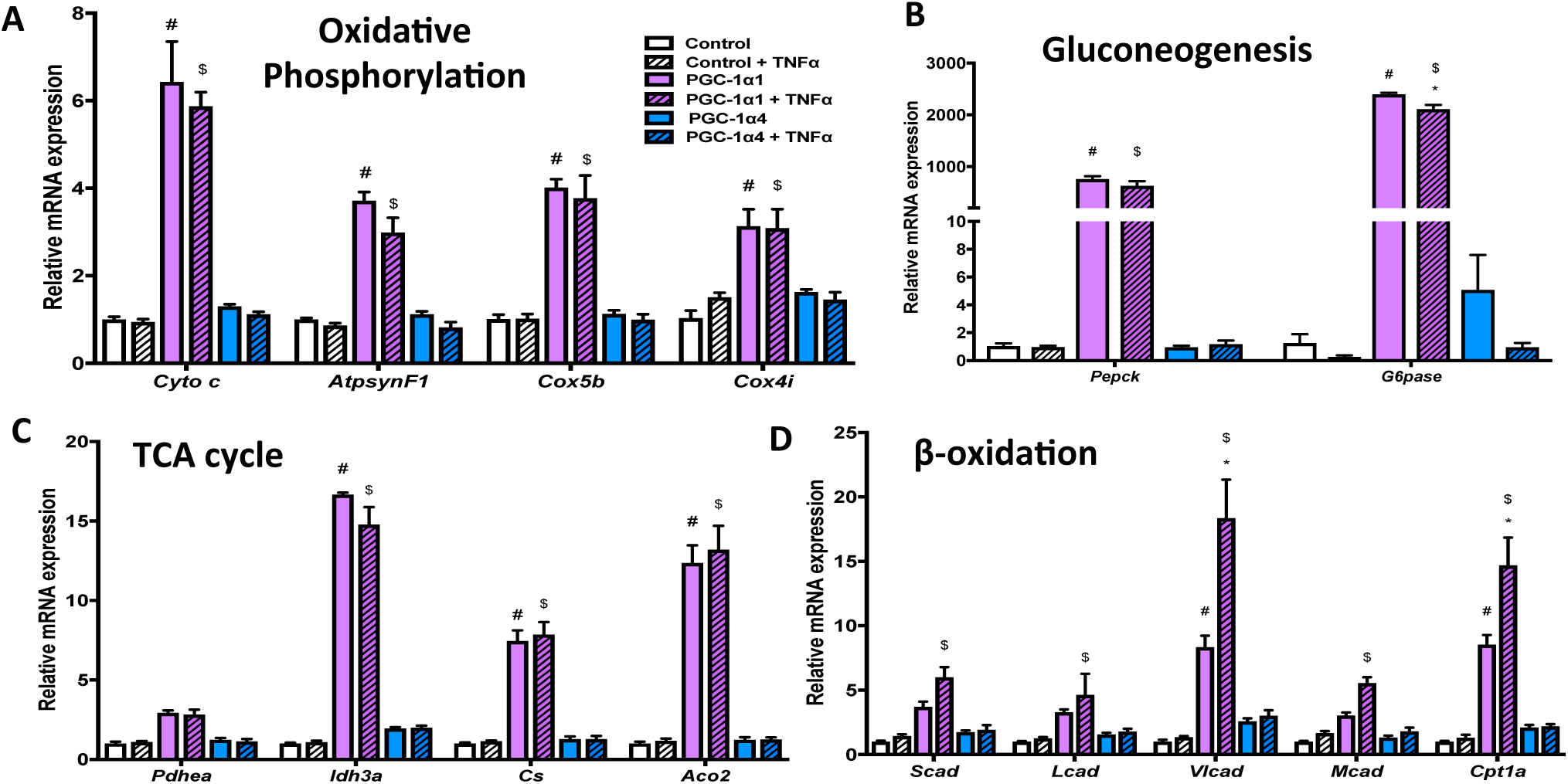
PGC-1α isoforms differentially regulate metabolic genes downstream of TNFα. A-D) mRNA expression of primary mouse hepatocytes over-expressing PGC-1α1, PGC-1α4 or vector alone following 2-hr treatment with 2 ng/mL TNFα or vehicle (n=3). *p<0.05 effect of TNFα within each genotype. **^#^**p<0.05 Effect of PGC-1α1 or PGC-1α4 expression compared to Control. **^$^**p<0.05 TNFα response compared to Control + TNFα. Data is representative biological replicates (n=3) from one experiment performed at least 3 independent times.

**Supplementary Figure 4:**
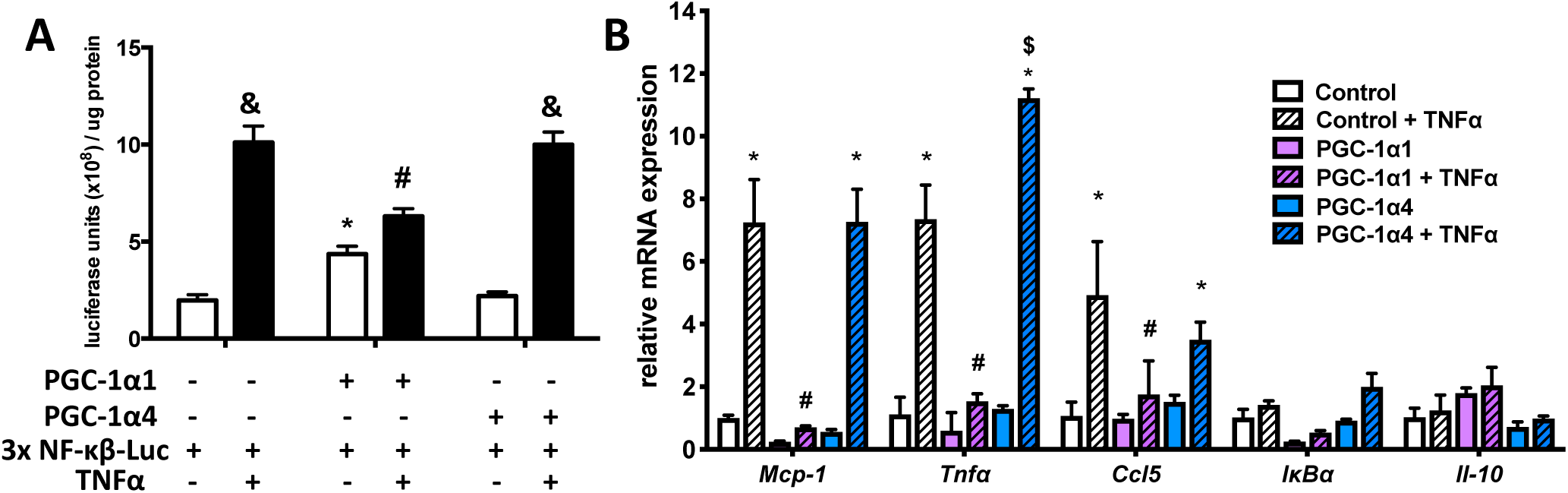
PGC-1α1, but not PGC-1a4 represses NF-κB activity and pro-inflammatory gene expression. A) Luciferase activity in primary mouse hepatocytes treated with 2 ng/ml TNFα or vehicle 48 hours following transfection with a 3x NF-κB reporter and constructs for PGC-1α1 or PGC-1α4 (or vector alone, n=3). *p<0.05 genotype effect compared to reporter vector, **^&^**p<0.05 TNFα response compared to vehicle, **^#^**p<0.05 TNFα response compared to vector + TNFα. B) mRNA expression of primary mouse hepatocytes over-expressing PGC-1α1, PGC-1α4 or vector following 2-hr treatment with 2 ng/mL TNFα or vehicle (n=3).

**Supplementary Figure 5:**
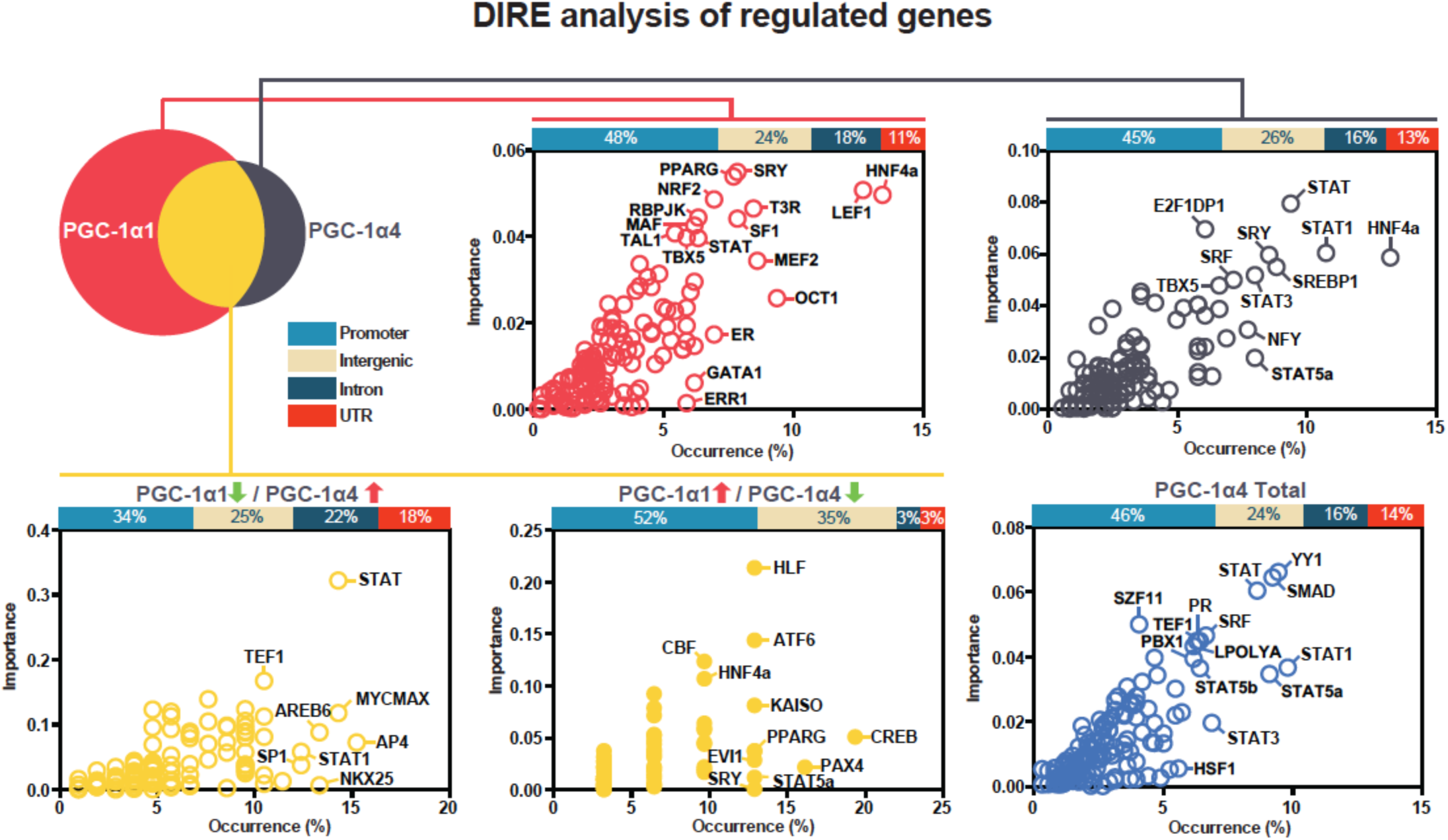
DiRE analysis of regulatory regions for enrichment of transcription factor binding sites. Ploeed are the occurrence of each binding motif and its importance metric, which reflects binding site specificity to the input gene set, compared to a background random set of 5000 genes. The top horizontal bars depict relative distribution of identified regulatory elements in promoters, intergenic, intronic, or untranslated regions.

**Supplementary Figure 6:**
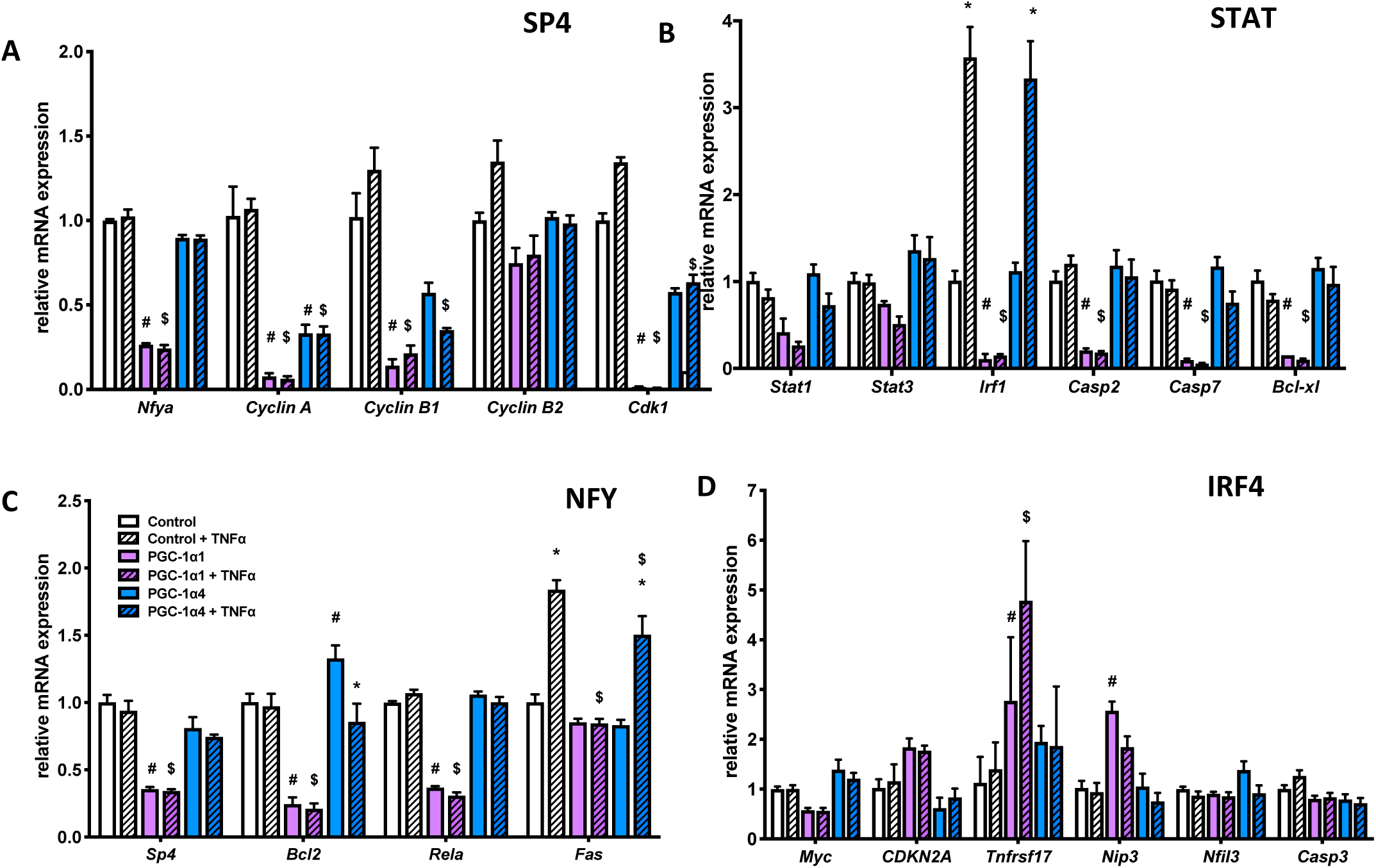
Target genes downstream of transcription factors identified by iRegulon and DiRE. A-D) mRNA expression of primary mouse hepatocytes over-expressing PGC-1α1, PGC-1α4 or vector alone following 2-hr treatment with 2 ng/mL TNFα or vehicle (n=3). *p<0.05 effect of TNFα within each genotype. **^#^**p<0.05 Effect of PGC-1α1 or PGC-1α4 expression compared to Control. **^$^**p<0.05 TNFα response compared to Control + TNFα. Data are representative of at least 2 different experiments. Data represents biological replicates (n=3) from a representative experiment performed twice.

**Supplementary Figure 7:**
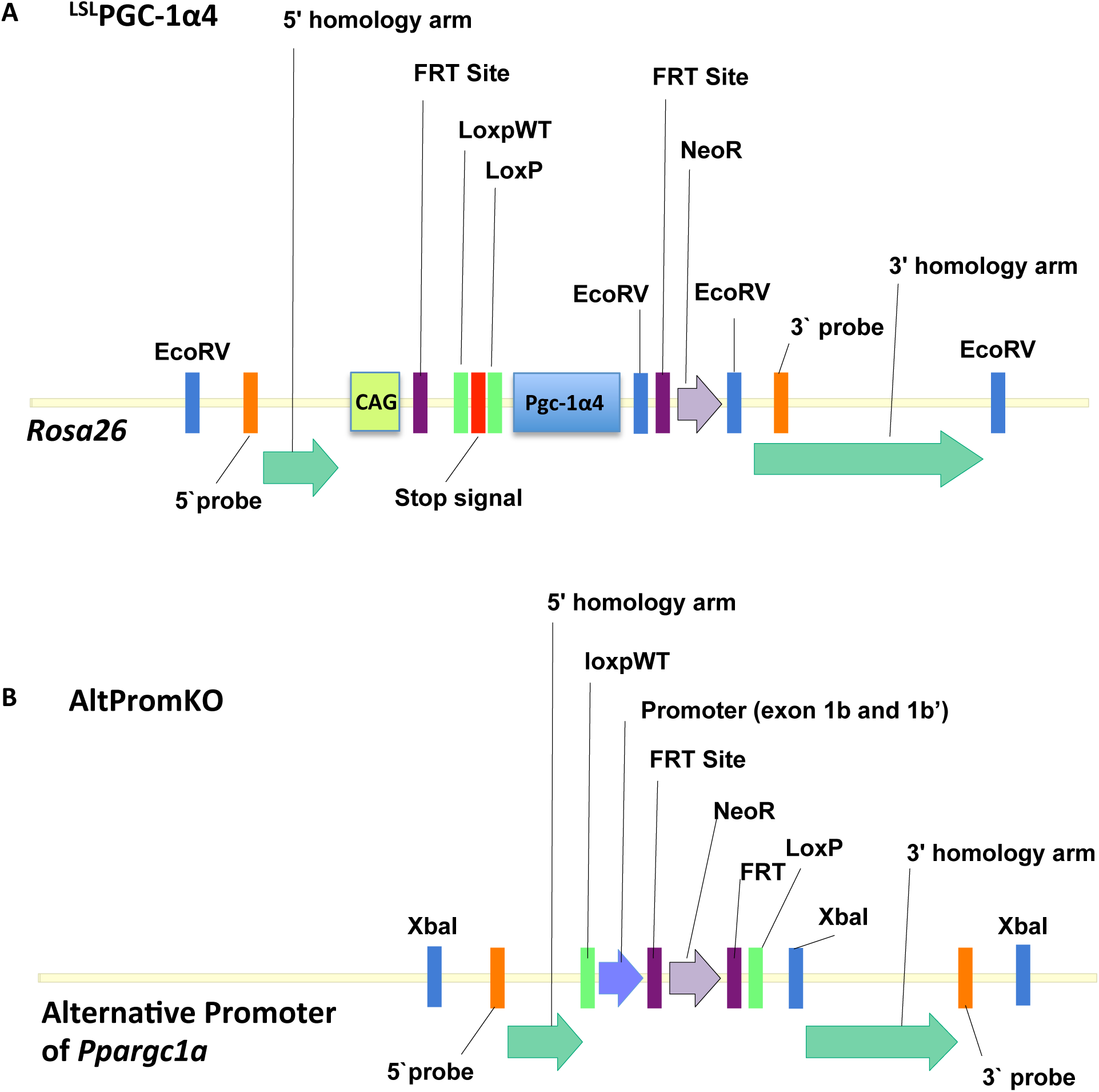
Targeting strategies for ^LSL^PGC-1α4 and AltPromKO mouse lines. Schematic of genomic locus for each mouse line, following recombination. Restriction sites (blue) and DNA probes (orange) used for Southern blot screening of ES clones are indicated. Complete diagrams of regulatory and cDNA elements inserted into genome can be found in Figures 4 and 7. A) The ^LSL^PGC-1α4 was generated by replacing tdTomato with murine PGC-1α4 cDNA in the Ai9 vector downstream of the Lox-stop-Lox signal. Recombination at the *ROSA26* locus was confirmed in neomycin-resistant C57BL/6 embryonic stem cells clones and founder mice backcrossed 10 generations onto a C57BL/6N background. B) The AltPromKO was generated by InGenious Targeting Laboratory (Ronkonkoma, New York). A targeting construct was used to insert LoxP sites flanking exon 1b and 1b’ of the alternative *Ppargc1a* promoter. Recombination was confirmed in C57BL/6 embryonic stem cells and founder mice backcrossed three times with C57BL/6N mice.

