## Supplementary Table 1 for "PGC-1α isoforms coordinate to balance hepatic metabolism and apoptosis in inflammatory environments"

| **Supplementary Table 1:** **Antibodies and Dilutions (**Target, company, catalog number, dilution) | | | |
| --- | --- | --- | --- |
| **Target Company Catalog Number Dilution** | | | |
| PGC-1α | Millipore | ST1202 | 1:500 |
| V5 | Thermo Scientific | MA5-15253 | 1:500 |
| Hsp90 | Cell Signaling | 4874 | 1:2000 |
| Cleaved Caspase 3 (Asp175) | Cell Signaling | 9661 | 1:500 |
| NFκB p50/p150 | Abcam | ab32360 | 1:500 |
| NFκB p65 | Abcam | ab7970 | 1:500 |
| IκBα | Abcam | ab32518 | 1:500 |
| IKKβ | Abcam | ab32135 | 1:1000 |
| Lamin B1 | BioVision | 3807 | 1:200 |
| β-actin | Sigma | A5441 | 1:5000 |
