## Supplementary Table 2 for "PGC-1α isoforms coordinate to balance hepatic metabolism and apoptosis in inflammatory environments"

| **Supplementary Table 2:** **Primers used for conventional PCR** (listed in 5’ – 3’ direction) | | | | | | |
| --- | --- | --- | --- | --- | --- | --- |
| Gene | | Forward Primer | | Reverse Primer | | Band size |
| **Mouse Primers** | | | | | | |
| *PGC-1α1* | | GACATGTGCAGCCAAGACTC | | CTCAAATGGGGAACCCTTGG | | 816 |
| *PGC-1α2* and *PGC-1α-b* | | GATTGTCATCCATGGATTC | | GTTCGCTCAATAGTCTTGTTC | | 325 / 826 |
| *PGC-1α3* and *PGC-1α-c* | | CTCAGACCCACTATGCTGCTG | | GTTCGCTCAATAGTCTTGTTC | | 302 / 818 |
| *PGC-1α4* | | GATTGTCATCCATGGATTC | | CTGGAAGATATGGCACAT | | 812 |
| *NT-PGC-1α-a* | | GACATGTGCAGCCAAGACTC | | CTGGAAGATATGGCACAT | | 822 |
| *NT-PGC-1α-c* | | CTCAGACCCACTATGCTGCTG | | CTGGAAGATATGGCACAT | | 803 |
| **Genotyping primers** | | | | | | |
| Alb-Cre^Tg^ | Forward (Albumin promoter)  TTAGAGGGGAACAGCTCCAGATGG | | Reverse (Cre-recombinase)  GTGAAACAGCATTGCTGTCACTT | |  | |
| ^LSL^PGC-1α4 | Forward (*Ppargc1a* exon 6)  CCAAACCAACAACTTTATCTC | | Reverse 1 (*Ppargc1a* intron 7) CCTTCTGATAAAGAGTCAACGC | | Reverse 2 (*WPRE*) GGAGAAAATGAAAGCCATACGG | |
| *Ppargc1a^fl/fl^* | Forward (*Ppargc1a* intron 2)  GGAGAGGTGTCAGGGAGAG | | Reverse (*Ppargc1a* intron 2)  CACAGCAGAGCACAAAGGA | |  | |
| AltProm*^fl/fl^* | Forward  AGAGTCAGCAGAACAAGCGT | | Reverse  TGCTTTGCAGAGGTGCTCAT | |  | |
