## Supplementary Table 3 for "PGC-1α isoforms coordinate to balance hepatic metabolism and apoptosis in inflammatory environments"

| **Supplementary Table 3:** Primers used for quantitative real-time PCR (listed in 5’ – 3’ direction) | | |
| --- | --- | --- |
| Gene | Forward Primer | Reverse Primer |
| **Mouse Primers** | | |
| *Birc2 (Ciap1)* | TCTGCTGTGGCCTGATGTTGGATA | ATGGAGACTGCAGACTGGCTGAAA |
| *Birc3 (Ciap2)* | AACTCCCTTCGGGAAATTGACCCT | TTCTTTCCTCCT GGAGTTTCCGCA |
| *Birc5 (survivin)* | TGGACAGACAGAGAGCCAAGAACA | AGCTGCTCAATTGACTGACGGGTA |
| *Ccl5 (Rantes)* | GCTGCTTTGCCTACCTCTCC | TCGAGTGACAAACACGACTGC |
| *IkBa (Nfkbia)* | AGACATCCTTCCGCAAACTC | TAGGTCCTTCCTGCCCATAA |
| *Il-10* | GCTCTTACTGACTGGCATGAG | CGCAGCTCTAGGAGCATGTG |
| *Mcp1 (Ccl2)* | TCACCTGCTGCTACTCATTCACCA | TACAGCTTCTTTGGGACACCTGCT |
| *Naip* | AGATGAAGAGCTCACCACCTGCTT | AGTTCAGTCAGTCTCATGGCAGCA |
| *Pgc-1α1/NT-PGC-1α-a* | GGACATGTGCAGCCAAGACTCT | CACTTCAATCCACCCAGAAAGCT |
| *Pgc-1α4/NT-PGC-1α-a,c* | TCACACCAAACCCACAGAAA | CTGGAAGATATGGCACAT |
| *Tnfα* | CCCTCACACTCAGATCATCTTCT | GCTACGACGTGGGCTACAG |
| *Tnfaip3 (A20)* | AGCCAGAAGAAGCTCAACTGGTGT | TGCATGCATGAGGCAGTTTCCATC |
| *Xiap* | CCAGCCATGGCAGAATATGA | TCGCCTTCACCTAAAGCATAAA |
| *Nfya* | CTCTGTGCCTGCTATCCAAA | CCTCTTAAGGATGCGGTGATAC |
| *Cyclin A* | CACTGACACCTCTTGACTATCC | CGTTCACTGGCTTGTCTTCTA |
| *Cyclin B1* | GGTCGTGAAGTGACTGGAAA | GTCTCCTGAAGCAGCCTAAAT |
| *Cyclin B2* | CTCTGCAAGATCGAGGACATAG | TGCCTGAGGTACTGGTAGAT |
| *Cdk1* | CAGACTTGAAAGCGAGGAAGA | TCCTGCAGGCTGACTATATTTG |
| *Cdc25c* | TGCACAGTCAGAAGGAACTG | GGAGGAGAATTCACAGAGGAAC |
| *Atad3a* | GACAGGACAGCACAGTAGTAAG | AGCAGACCATCTCGTCAATG |
| *Pim1* | TTCAGGCAAACGGTCTCTTC | CCACGGATGGTTCTGGATTT |
| *Csnk2a2* | CACATAGACCTAGATCCACACTTC | CAAGGTGCCTGTTCTCACTAT |
| *Btg2* | CGCACTGACCGATCATTACAA | GGGTCCATCTTGTGGTTGATAC |
| *Myc* | CTC CGT ACA GCC CTA TTT CAT C | TGG GAA GCA GCT CGA ATT T |
| *CDKN2A (p16)* | CAT GTT GTT GAG GCT AGA GAG G | CAC CGT AGT TGA GCA GAA GAG |
| *Tnfrsf17* | GCCTGGAGTATACAGTGGAAGA | CGGGAAGAAATGGTCAGAATCC |
| *Nip3* | GACGAAGTAGCTCCAAGAGTTC | CCAAAGCTGTGGCTGTCTAT |
| *Nfil3* | GGTTTCCGAAGCTGAGAATTTG | AGATCGGTTGTGTGGCTATG |
| *Casp3* | AGTGGGACTGATGAGGAGAT | GTAACCAGGTGCTGTAGAGTAAG |
| *Sp4* | TTTCTCAGCCAGCTTCTAGTTC | GGGTGGAAGGATTACCTGATTT |
| *Bcl2* | GGAGGATTGTGGCCTTCTTT | GTTCAGGTACTCAGTCATCCAC |
| *RelA (p65)* | GAGAAGCACAGATACCACCAAG | GAGATTCGAACTGTTCCTGGTC |
| *Fas* | CCAAGTGCAAGTGCAAACCAGACT | AGGATGGTCAACAACCATAGGCGA |
